# Tozorakimab (MEDI3506): a dual-pharmacology anti-IL-33 antibody that inhibits IL-33 signalling via ST2 and RAGE/EGFR to reduce inflammation and epithelial dysfunction

**DOI:** 10.1101/2023.02.28.527262

**Authors:** Elizabeth England, D. Gareth Rees, Ian Christopher Scott, Sara Carmen, Denice T. Y. Chan, Catherine E. Chaillan Huntington, Kirsty F. Houslay, Teodor Erngren, Mark Penney, Jayesh B. Majithiya, Laura Rapley, Dorothy A. Sims, Claire Hollins, Elizabeth C. Hinchy, Martin D. Strain, Benjamin P. Kemp, Dominic J. Corkill, Richard D. May, Katherine A. Vousden, Robin J. Butler, Tomas Mustelin, Tristan J. Vaughan, David C. Lowe, Caroline Colley, E. Suzanne Cohen

## Abstract

Interleukin (IL)-33 is a broad-acting alarmin cytokine that can drive inflammatory responses following tissue damage or infection and is a promising target for treatment of inflammatory disease. Here, we describe the identification of tozorakimab (MEDI3506), a potent, human anti-IL-33 monoclonal antibody, which can inhibit reduced IL-33 (IL-33^red^) and oxidized IL-33 (IL-33^ox^) activities through distinct serum-stimulated 2 (ST2) and receptor for advanced glycation end products - epidermal growth factor receptor (RAGE-EGFR complex) signalling pathways. We hypothesized that a therapeutic antibody would require an affinity higher than that of ST2 for IL-33, with an association rate greater than 10^7^ M^−1^ s^−1^, to effectively neutralize IL-33 following rapid release from damaged tissue. An innovative antibody generation campaign identified tozorakimab, an antibody with a femtomolar affinity for IL-33^red^ and a fast association rate (8.5 × 10^7^ M^−1^ s^−1^), which was comparable to soluble ST2. Tozorakimab potently inhibited ST2-dependent inflammatory responses driven by IL-33 in primary human cells and in a murine model of lung epithelial injury. Additionally, tozorakimab prevented the oxidation of IL-33 and its activity via the RAGE/EGFR signalling pathway, thus increasing *in vitro* epithelial cell migration and repair. Tozorakimab is a novel therapeutic agent with a dual mechanism of action that blocks IL-33^red^ and IL-33^ox^ signalling, offering potential to reduce inflammation and epithelial dysfunction in human disease.

## Introduction

Interleukin (IL)-33 is a broad-acting IL-1 family cytokine that is released from stressed or damaged barrier tissues, including the endothelium and epithelium, following external triggers such as trauma, allergen exposure or infection [1,2]. Under physiological conditions, IL-33 initiates protective immune responses; however, excess IL-33 release or chronic signalling can drive tissue-damaging inflammation and remodelling [1,3,4]. Almost two decades of pre-clinical evidence suggests that dysregulated IL-33 activities may contribute to the pathology of inflammatory diseases and severe infectious diseases, including COVID-19 [1,2,5–10]. This is further supported by clinical efficacy data for antibodies to IL-33 and its receptor serum-stimulated 2 (ST2; also named IL1RL1) that have provided clinical precedence for targeting IL-33 in chronic obstructive pulmonary disease (COPD) and asthma [11–14].

IL-33 is localized to the nucleus via N-terminal sequences and chromatin-binding domains [15]. The full-length IL-33 protein is biologically active; however, its activity through ST2 is enhanced up to 60-fold by the removal of the N-terminal domain [16–20]. IL-33 exists in both reduced (IL-33^red^) and oxidized (IL-33^ox^) forms that signal via distinct downstream pathways [21,22].

IL-33^red^ is a member of the IL-1 receptor family and signals via ST2 [23]; ST2 is expressed as two isoforms: a membrane-associated variant (ST2L) and a truncated, soluble form (sST2). The truncated, soluble form lacks the transmembrane and intracellular domains of ST2L [24,25].

IL-33^red^ exerts cellular functions through the receptor complex of ST2L and the IL-1 receptor accessory protein [24]. ST2 is constitutively expressed on some immune (e.g. mast cells and type 2 innate lymphoid cells) [26] and endothelial cells and can be induced (e.g. by IL-12) on additional immune cell types such as natural killer cells [7]. On binding to ST2, IL-33^red^ initiates nuclear factor kappa-light-chain-enhancer of activated B cells (NF-κB) and mitogen-activated protein kinase signalling [1,27]. This results in a cascade of pro-inflammatory signalling pathways, including the release of cytokines and chemokines [1,3,28,29]. IL-33 activity is regulated by sST2, which is an endogenous antagonist of IL-33 [30].

IL-33^ox^ cannot signal via ST2 [21]. Oxidation was initially suggested by our group as a mechanism of inactivation of IL-33 [21]. However, our subsequent studies have shown that human IL-33^ox^ binds to the receptor for advanced glycation end products (RAGE) and signals via a complex with the epidermal growth factor receptor (EGFR) [22]. The IL-33^ox^ RAGE-EGFR signalling pathway can drive remodelling of the airway epithelium, resulting in mucus hypersecretion in an *in vitro* model of COPD [22].

Targeting the IL-33-ST2 axis is a therapeutic strategy under clinical investigation for inflammatory diseases [11,31–34]. Here, we describe tozorakimab (MEDI3506), a novel high-affinity anti-IL-33 human monoclonal antibody generated via an innovative lead generation campaign using an oxidation-resistant form of recombinant IL-33. To the best of our knowledge, tozorakimab is the first anti-IL-33 antibody described that inhibits the activity of both IL-33^red^ and IL-33^ox^ through the ST2 and RAGE-EGFR signalling pathways, respectively.

## Results

### An effective anti-IL-33 antibody is predicted to require a high affinity and a fast association rate

To gain an understanding of the binding kinetics and affinity required for a therapeutically effective anti-IL-33 antibody, we first determined the binding kinetics of IL-33 to sST2. The affinity of IL-33^red^ for sST2 was 0.09 pM (95% confidence interval [CI], 0.05–0.15; Fig. 1a–c), which was determined using a highly sensitive kinetic exclusion assay (KinExA) [35]. Active constant binding partner values indicated that approximately 65% of sST2 was active and/or participating in the interaction. IL-33^red^ bound to sST2 with a fast association rate (1.5 × 10^8^ M^−1^ s^−1^; 95% CI, 1.4–1.6 × 10^8^; Fig. 1d and e).

**Figure 1.**
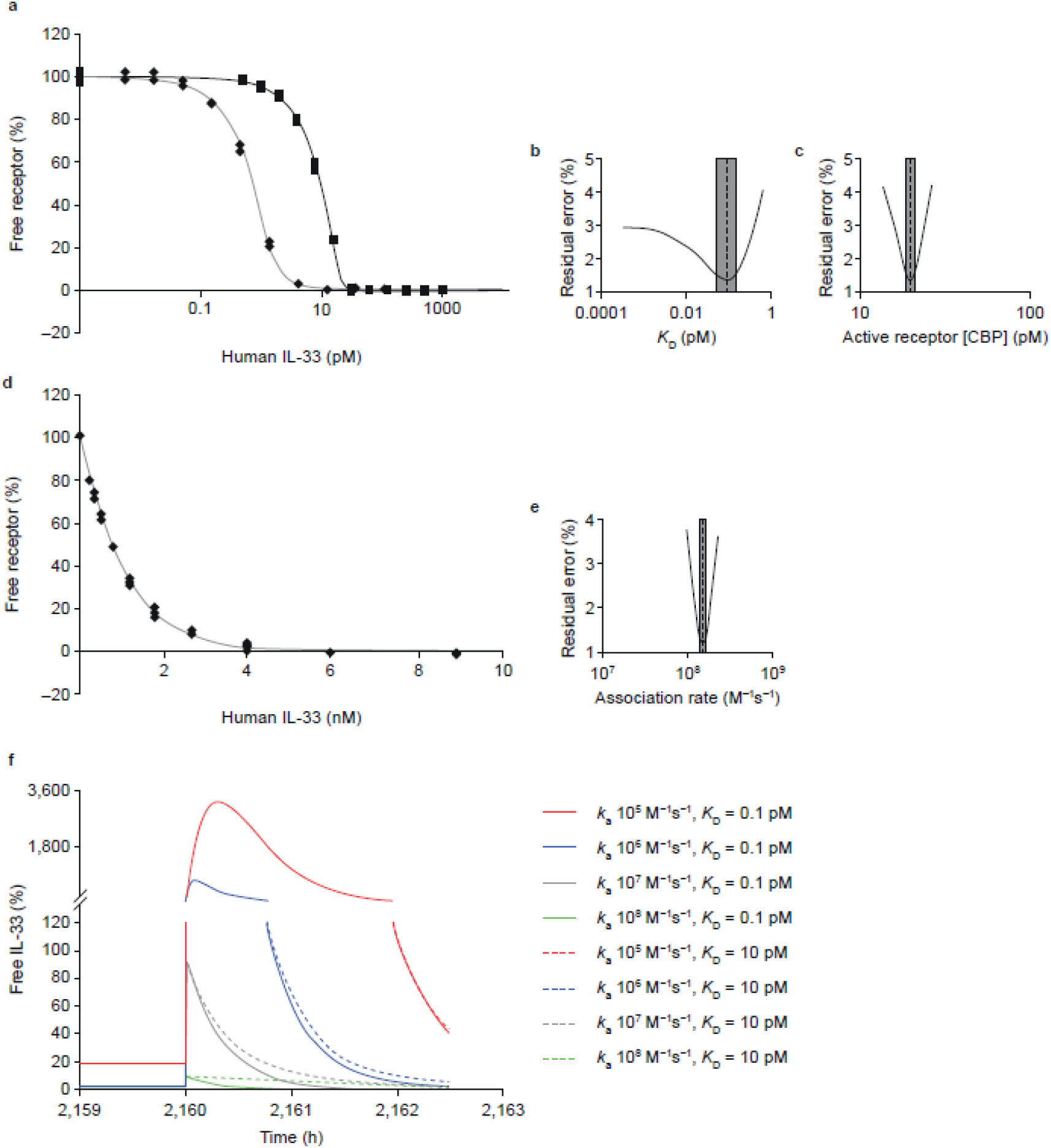
IL-33 binds sST2 with a high affinity and a fast association rate. **(a)** The affinity (*K*_D_) of IL-33^red^ (residues 112–270) interacting with sST2 was obtained after a 1:1 binding model was fitted to two KinExA datasets simultaneously; the black solid line with filled squares are the 30 pM fixed [sST2] data (■) and the grey line with filled diamonds are the 2.0 pM fixed [sST2] data (⍰). **(b)** A plot of 95% CIs (indicated by the grey rectangle) around the affinity estimate (dashed line) shown in panel a. **(c)** A plot of 95% CIs around the active receptor CBP parameter shown in panel a. **(d)** The association rate constant (*k*_a_) for IL-33 interacting with sST2 using KinExA. **(e)** The 95% CI around the association rate constant estimate shown in panel d. **(f)** In silico modelling of the suppression profile expected from antibodies with varying kinetic and affinity profiles, during acute IL-33 spike profiles parameterized using measured IL-33 spike data from ALT challenged mice, following the establishment of steady state suppression. The spike suppression was modelled assuming prior dosing of 150 mg neutralizing antibody every 4 weeks with an IL-33 degradation half-life of 60 minutes and an IL-33 spike half-life of 15 min (spike occurring at 2,160 hours; refer to supplementary Figure 1a). Antibody affinity (*K*_D_) was varied over a 0.1–10 pM range (solid versus dashed lines, respectively) with the association rate (*k*_a_), which was varied in 10-fold steps over the 10^5^ to 10^8^ M^−1^ s^−1^ range (red, blue, grey and green, respectively). *ALT, Alternaria alternata*; CBP, constant-binding partner; CI, confidence interval; IL, interleukin; KinExA, Kinetic Exclusion Assays; sST2, soluble serum stimulated 2.

Based on the observed affinity of IL-33^red^ for sST2, it was hypothesized that a high-affinity anti-IL-33 antibody would be required to neutralize IL-33 activity. The binding kinetics data generated above were used to inform an *in silico* analysis that was performed to understand antibody performance. Given that rapid release of IL-33 into the lung lumen has previously been observed during epithelial injury [16,17], antibody effects were also modelled on the suppression of spikes of increased IL-33 release associated with disease exacerbation [36] (**Supplementary Fig. S1**).

When the anti-IL-33 antibody affinity was narrowed to encompass a range of very high affinities between 0.1 and 10 pM in the *in silico* model, the effects of affinity gain on predicted free IL-33^red^ were modest (**Fig. 1f**). However, the *in silico* model did predict that the anti-IL-33 antibody association rate with IL-33 was a key driver of IL-33 suppression. Very fast association rates, of between 10^7^ and 10^8^ M^−1^ s^−1^, attenuated the free IL-33 spike below 100 and 10% of steady-state levels, respectively (**Fig. 1f**).

Based on the *in silico* modelling results, it was hypothesized that a high-affinity antibody with a fast association rate would be needed to effectively neutralize IL-33 that is rapidly released during tissue injury.

### Identification of an anti-IL-33 antibody that neutralizes IL-33^red^ and ST2-dependent activities

A novel discovery campaign was devised to identify antibodies that selectively bind IL-33^red^ and prevent its interaction with ST2. As IL-33^red^ is susceptible to rapid oxidation and associated conformational changes [21], we aimed to preserve the conformational epitopes present on IL-33^red^ by replacing cysteine with serine residues to produce oxidation-resistant IL-33 (IL-33^C>S^) [21]. IL-33^C>S^ has ST2-dependent activity like that of IL-33^red^ and is protected from a loss of ST2-dependent activities due to oxidation [21].

Recombinant IL-33^C>S^ and IL-33^red^ (both comprising IL-33 residues 112–270) were used to identify anti-IL-33 antibodies. Both forms of IL-33 were presented to naive phage libraries [37–39] expressing single-chain variable fragments (scFvs). Binding to IL-33^C>S^ or IL-33^red^ was enriched during three rounds of selection. Over 8500 scFvs were screened for IL-33 neutralization using IL-33^C>S^–sST2 and IL-33^red^–sST2 competition assays. In total, 71 IL-33^red^ neutralizing scFvs were identified, 41 from IL-33^C>S^ and 30 from IL-33^red^, which were converted to human immunoglobulin G (IgG) 1.

The functional activities of the anti-IL-33 neutralizing antibodies were studied in human umbilical vein endothelial cells (HUVECs) [21]. It was hypothesized that an anti-IL-33 antibody would inhibit both NF-κB signalling and cytokine release in order to be therapeutically effective. A single lead monoclonal antibody (mAb; clone number, 33v20064) which inhibited both NF-κB signalling and cytokine release was identified. The lead mAb was derived from selections using IL-33^C>S^ and inhibited IL-33–sST2 binding in a biochemical competition assay (**Fig. 2a**), NF-κB signalling and IL-8 release in HUVECs (**Fig. 2b and c**). However, the lead mAb had a modest affinity for IL-33^red^ (29 nM) in surface plasmon resonance studies (**Fig. 2d**). In summary, these data indicated that significant affinity optimization was required to produce an anti-IL-33 antibody with the potential to be therapeutically effective.

**Figure 2.**
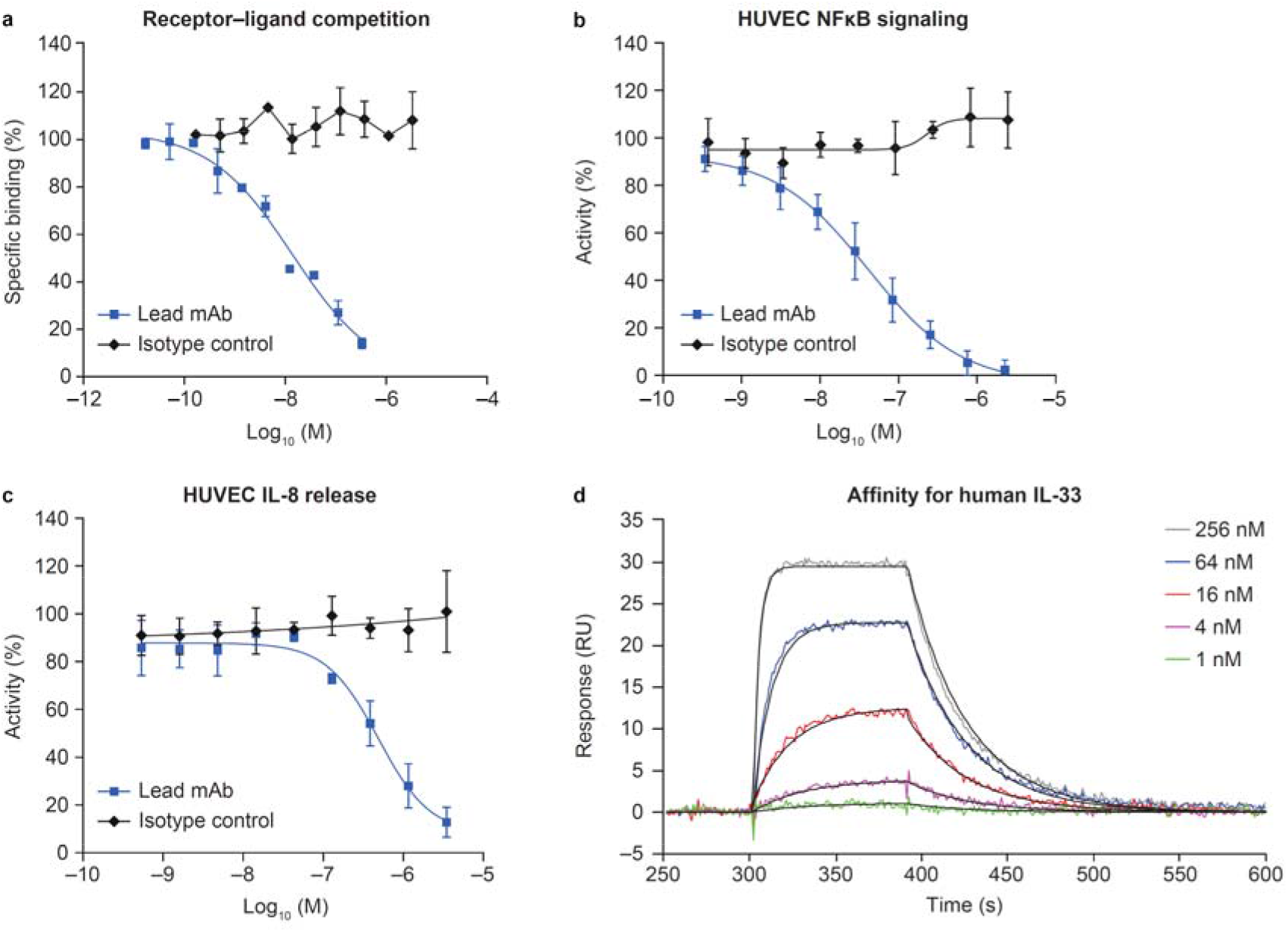
Characterization of the lead antibody (33v20064) in biochemical and *in vitro* assays. **(a)** IL-33 binding sST2 biochemical competition assay (mean ± SD of duplicate determinations). **(b)** Inhibition of NF-κB signalling in HUVECs (mean ± SD, n = 3) **(c)** Inhibition of IL-8 release in HUVECs (mean ± SD, n = 2). **(d)** Affinity of lead antibody for human IL-33 at concentrations of 1–256 nM, which was determined by surface plasmon resonance. For each sensorgram trace 1:1 fit lines are shown in black. Figure shows a representative dataset from four separate experiments. HUVEC, human umbilical vein endothelial cell; IL, interleukin; mAb, monoclonal antibody; NF-κB, nuclear factor kappa-light-chain-enhancer of activated B cells; RU, response units; SD, standard deviation.

### Generation of tozorakimab, a potent, high-affinity IL-33 neutralizing antibody

The affinity of the lead mAb (33v20064) for IL-33^red^ was at least 100 000-fold lower than the affinity of sST2 (< 90 fM) for IL-33^red^. Therefore, comprehensive affinity maturation was undertaken via random mutagenesis of all six complementarity-determining regions (CDRs) [40,41]. Phage libraries of the lead mAb (33v20064) were generated and scFv variants were enriched for binding to IL-33^C>S^. Variants with an improved affinity were identified by screening using an IL-33^C>S^–lead mAb (33v20064) epitope competition assay. Beneficial mutations identified in the variable fragment heavy chain (VH) and variable fragment light chain (VL) CDRs were recombined; additional changes in framework regions were made to match the closest human germline sequences to generate tozorakimab. In total, tozorakimab had five amino acid changes in VH CDR2, four changes in VH CDR3, one change in VL CDR1, one change in VL CDR2, five changes in VL CDR3, and required six amino acid changes in the VL framework to match the closest human germline sequence from the starting lead mAb (33v20064).

Tozorakimab had a >10 000-fold higher potency for IL-33^red^ than the lead mAb (33v20064) in a biochemical epitope competition assay (half maximal inhibitory concentration [IC_50_]: tozorakimab, 0.082 nM; lead mAb, 689 nM; **Fig. 3a**). Tozorakimab also completely inhibited IL-33 binding to sST2 in an IL-33–sST2 competition assay (**Fig. 3b**) and inhibited recombinant IL-33^red^ driven IL-8 release (**Fig. 3c**) in HUVECs with a potency approximately 10-fold greater than sST2 (Table 1). Tozorakimab also demonstrated potent inhibition of IL-33-driven inflammatory mediators (IL-13, IL-6, IL-8, granulocyte-macrophage colony-stimulating factor and tumour necrosis factor alpha) in human blood-derived mast cells (**Fig. 3d and Table 1; Supplementary Fig. S2a–e**,) and interferon-gamma (IFN-γ) release from peripheral blood mononuclear cells (PBMCs; **Fig. 3e and Table 1**).

**Table 1.**
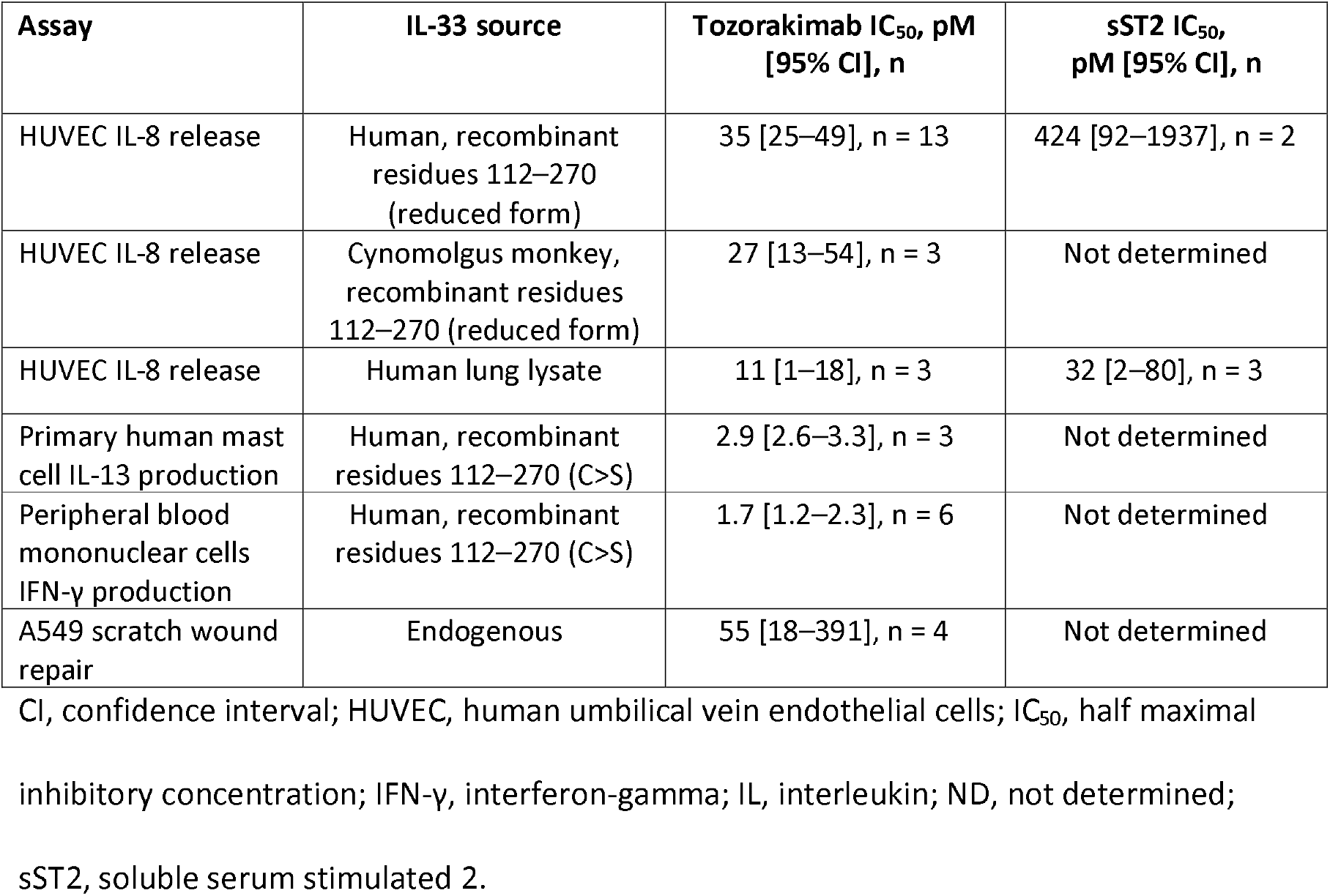
Summary of potencies of tozorakimab *in vitro*.

**Figure 3.**
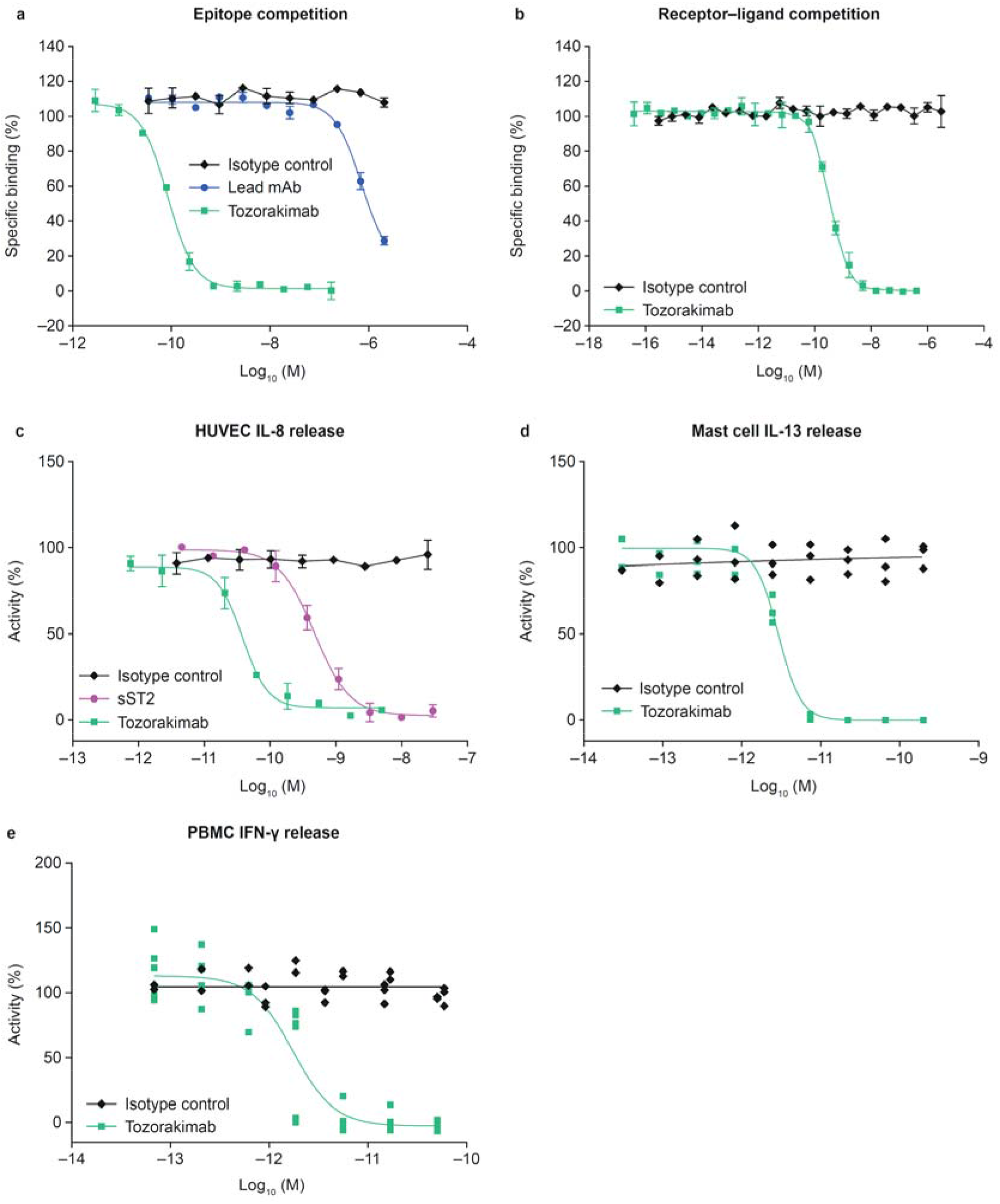
*In vitro* pharmacology of tozorakimab. **(a)** Epitope competition assay. **(b)** Biochemical receptor–ligand competition assay. **(c)** Inhibition of recombinant IL-33^red^ (residues 112–270)-driven IL-8 release in HUVECs. **(d)** Inhibition of recombinant IL-33^C>S^ (residues 112–270)-driven IL-13 release in human blood-derived mast cells (three individual donors shown, data points are the mean of duplicate tests). **(e)** Inhibition of recombinant IL-33^C>S^ (residues 112–270)-driven IFN-γ release from human PBMCs (six individual donors, data points are the mean of duplicate tests). **(a–c)**. The data shown are a representative example from at least two independent experiments. Data points are plotted as mean ± SD of duplicate determinations. HUVEC, human umbilical vein endothelial cell; IFN-γ, interferon-gamma; IL, interleukin; PBMC, peripheral blood mononuclear cell; red, reduced; SD, standard deviation; sST2, soluble serum stimulated 2.

Tozorakimab did not bind to closely related human IL-1 family proteins IL-1 alpha and IL-1 beta, which confirmed its specificity for IL-33 (**Supplementary Fig. S3a**). To assess the potential for safety and pharmacology studies to be undertaken in animals, cross-reactivity to cynomolgus monkey IL-33 was confirmed using a biochemical epitope competition (IC_50_, 11.07 nM; 95% CI, 9.62–12.73; **Supplementary Fig. S3a**) and HUVEC IL-8 release assays (**Table 1; Supplementary Fig. S3b and c**). However, tozorakimab did not bind rodent IL-33 (**Supplementary Fig. S3a**).

### Tozorakimab has a fast association rate and superior affinity for IL-33^red^

The affinity of tozorakimab for IL-33^red^ was 3-fold higher than the affinity of sST2 for IL-33^red^ (30 vs 90 fM; **Fig. 1a and 4a–c; Supplementary Table S1**), which was consistent with the relative *in vitro* potencies for tozorakimab and sST2. A monovalent antigen-binding fragment (Fab) of tozorakimab retained a similar high affinity for IL-33^red^, which confirmed that bivalent avid interactions did not significantly contribute to the affinity of tozorakimab (**Supplementary Table S1**). Tozorakimab had a fast association rate when compared with sST2 (ka, 8.5 × 10^7^ vs 1.5 × 10^8^ M^−1^ s^−1^, respectively; **Fig. 4d and e; Supplementary Table S1**).

**Figure 4.**
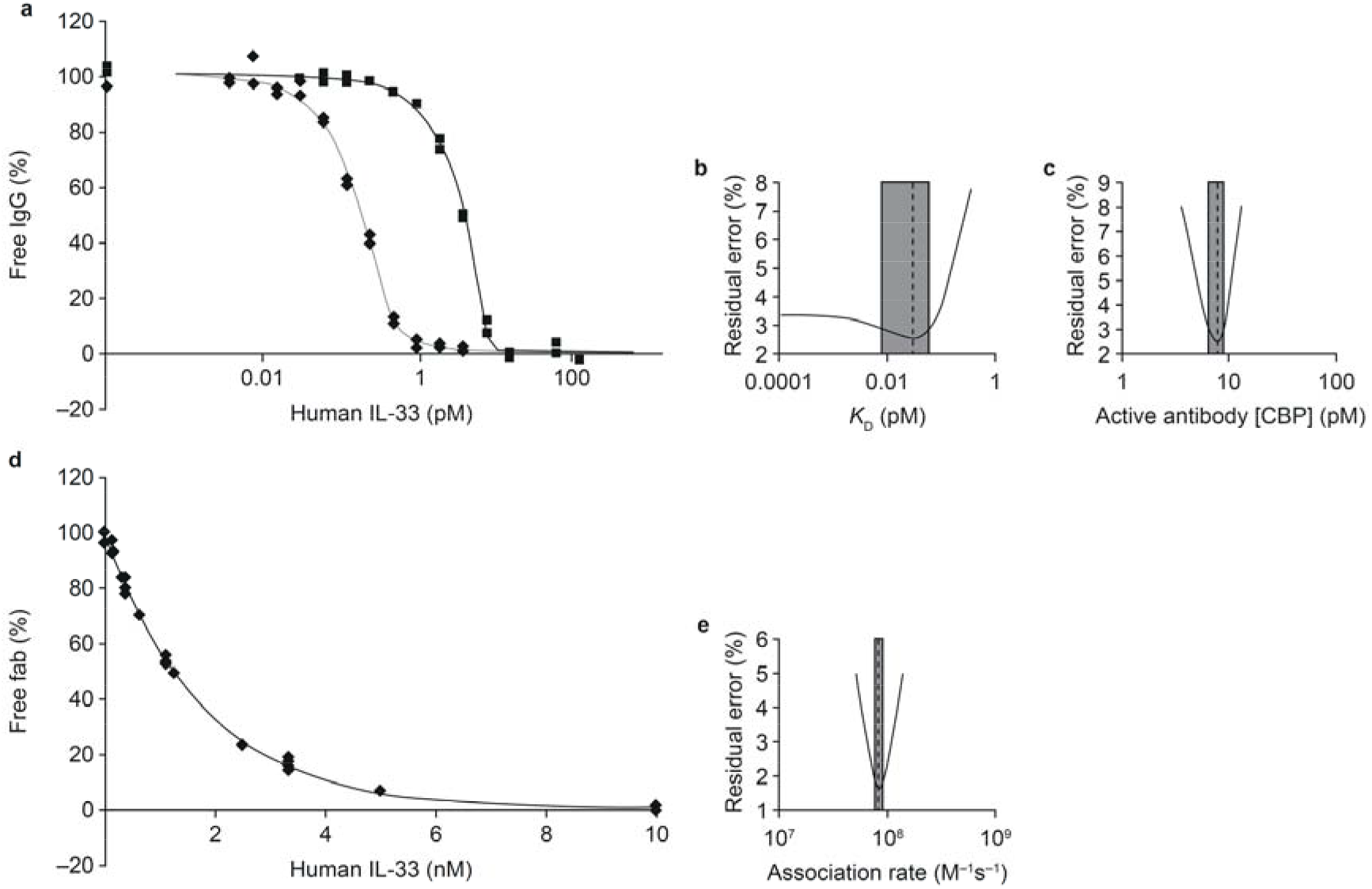
Affinity and association rate of tozorakimab for IL-33^red^ measured using KinExA. **(a)** The affinity (*K*_D_) of tozorakimab for IL-33^red^ (residues 112–270) was obtained after a 1:1 binding model was fitted to two KinExA datasets simultaneously; the black solid line and filled squares are the data from IL-33^red^ titrated into 5 pM fixed [IgG] (■) and the grey solid line and filled diamonds are from a 0.20 pM fixed [IgG] data titration (⍰). **(b)** Error curve indicating the extent (grey shaded rectangle) of 95% CIs around the affinity value (indicated by dashed line) from the fit to the data in panel a. **(c)** Fitting error plot of the 95% CIs of the active receptor CBP parameter that arise from the fit in panel a. **(d)** The association rate constant for the tozorakimab Fab fragment binding IL-33^red^ (residues 112–270) measured using KinExA. **(e)** Error plot of the 95% CI around the association rate (k_a_) estimate from the fit in panel d. CBP, constant-binding partner; CI, confidence interval; Fab, antigen-binding fragment; IgG, immunoglobulin G; KinExA, kinetic exclusion assays; IL, interleukin; red, reduced.

### Tozorakimab inhibits full length and N-terminally processed forms of endogenous IL-33^red^

Full length and N-terminally processed forms of endogenous IL-33^red^ have activity at the ST2 receptor [18]. Tozorakimab potently inhibited a mixture of full-length and N-terminally truncated forms of IL-33^red^, derived from human lung tissue, with a potency approximately 3-fold greater than sST2 (**Fig. 5a and b and Table 1**).

**Figure 5.**
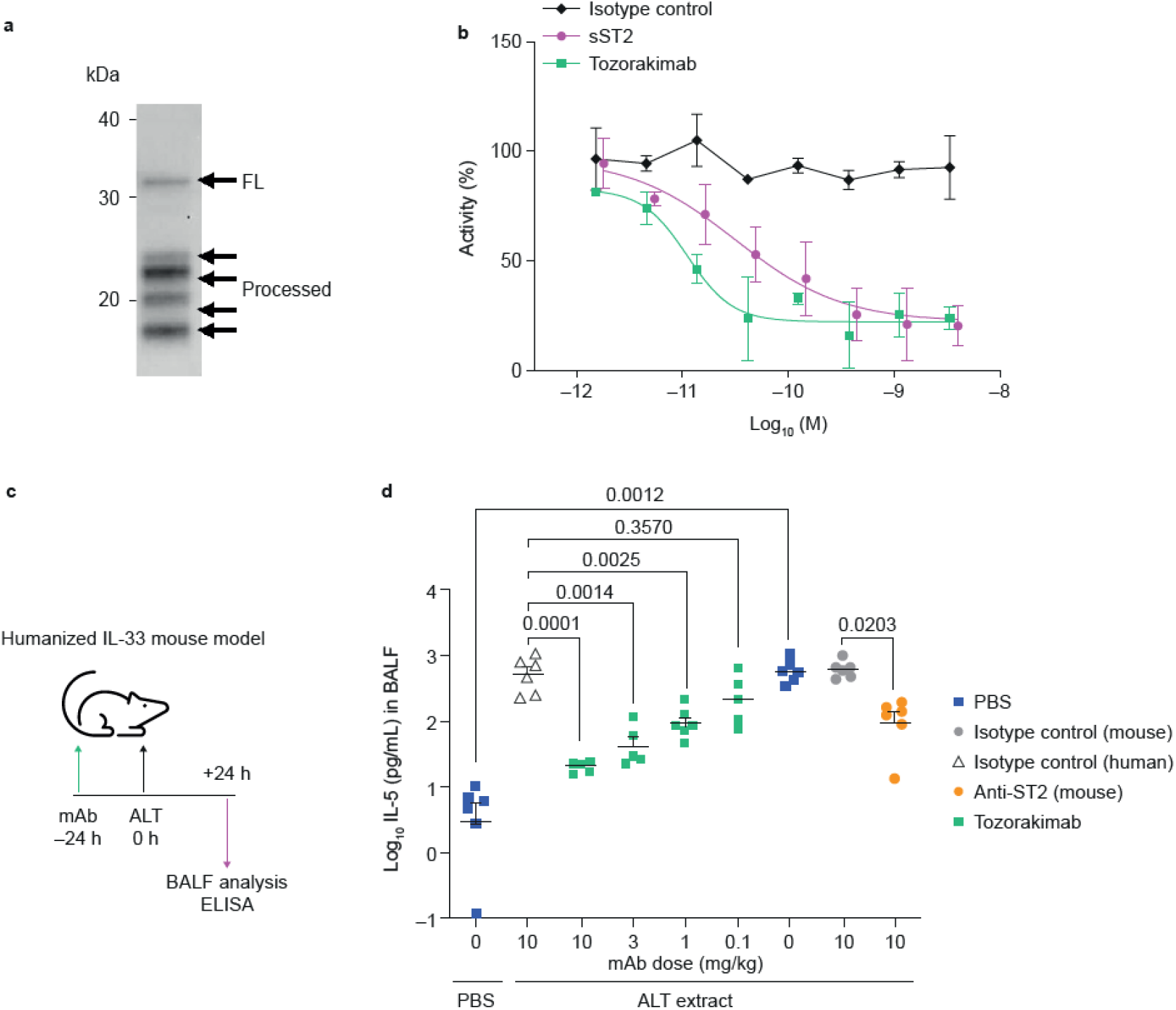
Tozorakimab can inhibit endogenous IL-33^red^-driven inflammatory responses *in vitro* and *in vivo*. **(a)** Western blot of human IL-33 in lung tissue lysate derived from a healthy individual who was an ex-smoker (representative of n = 3). **(b)** Inhibition of IL-8 release in HUVECs treated with human lung lysate (mean ± SD, n = 3). **(c)** Study to determine inhibition of endogenous IL-33-dependent IL-5 in BALF following an ALT challenge. Tozorakimab administered i.p. 24 h before ALT challenge. The study was performed in female humanized-IL-33 KI, mouse-IL-33 KO mice (huIL-33KI). **(d)** Inhibition of ALT driven IL-5 release in BALF. PBS, human-isotype mAb and mouse-isotype Mab controls were used (n = 6). A Browne–Forsyth ANOVA was used to test the impact of dose on log_10_ transformed IL-5 levels to confirm that the samples were of similar variance. The post hoc Dunnett’s multiple comparison test was then used to generate *p* values (as stated) for the comparisons of interest (Prism 8). ALT, *Alternaria alternata*; ANOVA, analysis of variance; BALF, bronchoalveolar lavage fluid; ELISA, enzyme-linked immunosorbent assay; FL, full length; HUVEC, human umbilical vein endothelial cell; IL, interleukin; i.p., intraperitoneal; KI, knock in; KO, knockout; mAb, monoclonal antibody; PBS, phosphate buffered saline; red, reduced; SD, standard deviation; sST2, soluble serum stimulated 2.

### Tozorakimab inhibits IL-33 activity *in vivo*

Humanized IL-33 (human IL-33^+/+^; mouse IL-33^−/−^) genetically modified mice were intranasally challenged with *Alternaria alternata* to induce lung epithelial injury and IL-33-dependent inflammation (**Fig. 5c**) [18,21,42]. Prophylactic intraperitoneal administration of tozorakimab (0.1, 1.0, 3.0 and 10.0 mg/kg) dose-dependently inhibited IL-5 levels in bronchoalveolar lavage fluid (BALF) compared with the controls (**Fig. 5d**). Anti-mouse ST2 mAb also inhibited BALF IL-5 in this model (**Fig. 5d)**, which was consistent with the hypothesis that tozorakimab inhibits ST2⍰dependent responses *in vivo*.

### Tozorakimab prevents oxidization of IL-33 and downstream IL-33 ox signalling via the RAGE/EGFR receptor complex

It was recently reported that, independently of ST2, human IL-33^ox^ binds RAGE and signals via a complex with EGFR. IL-33^ox^ RAGE EGFR signalling in airway epithelium basal cells impairs scratch wound healing responses [22]. To investigate tozorakimab inhibition of the IL-33^ox^ RAGE EGFR signalling pathway, its impact on IL-33^ox^-mediated biology was evaluated using a scratch wound healing assay in A549 epithelial cells that are reported to lack ST2 [25]. Tozorakimab enhanced A549 cell scratch wound healing (**Fig. 6a**), indicating potent neutralisation of IL-33^ox^ signalling (**Table 1**).

**Figure 6.**
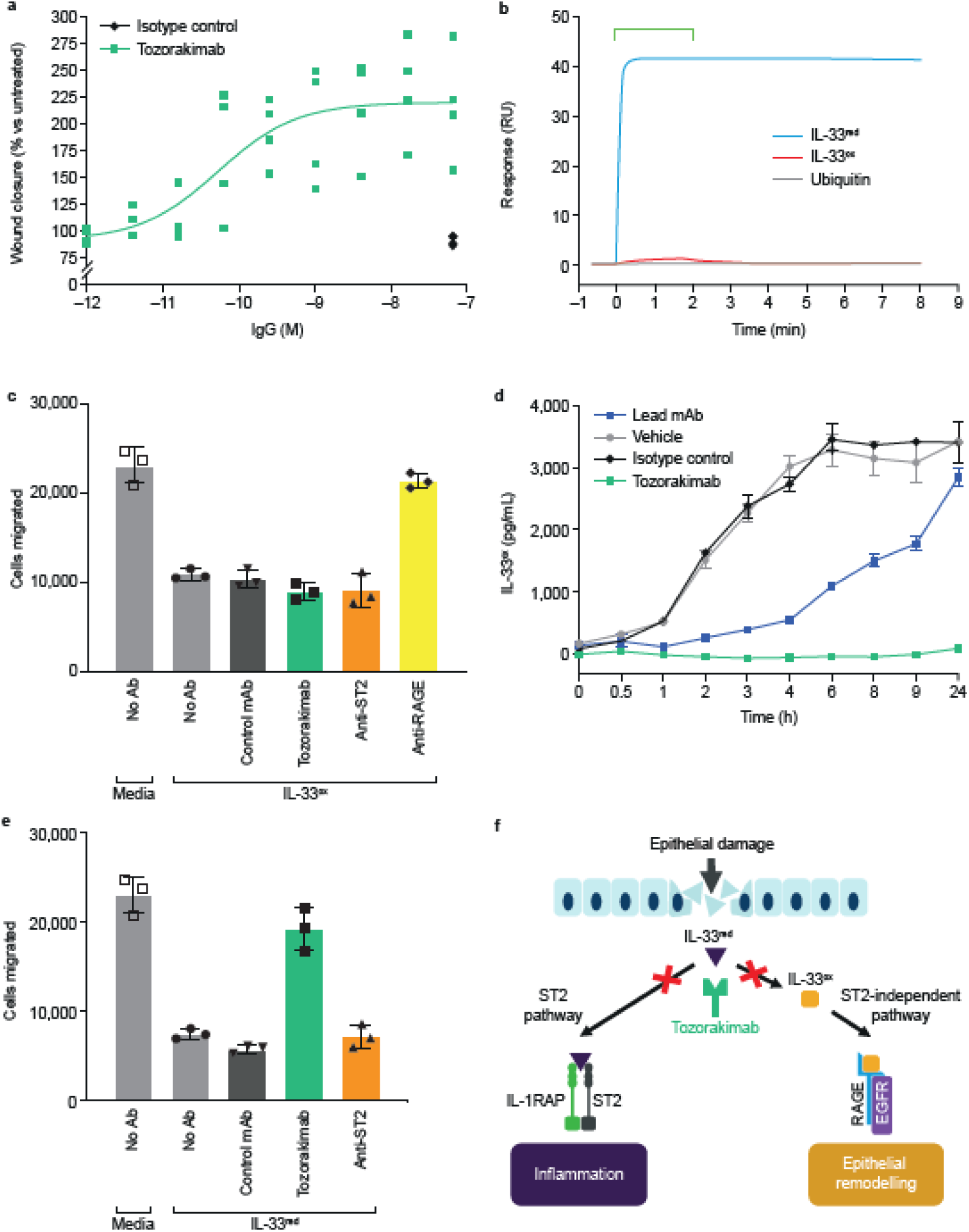
Tozorakimab potently inhibits the formation of IL-33^ox^. **(a)** The A549 epithelial cell scratch wound healing assay was normalized to the untreated control (data points are a mean of six replicates, n = 4 separate experiments). **(b)** Surface plasmon resonance binding profiles for tozorakimab Fab fragment binding to chip-immobilized N-terminally tagged wild-type IL-33^red^ or wild-type IL-33^ox^. Both IL-33 constructs were based on the 112–270 residue form. Amine biotinylated ubiquitin served as a protein control. The bar indicates the timing of the 2 min injection of 10 nM monomerized Fab. **(c)** Impact of recombinant IL-33^ox^ in the presence of test antibodies on human lung epithelial cell migration. The data shown (mean ± SD of triplicate determinations) are a representative example from two independent experiments. **(d)** Time course of recombinant IL-33^red^ oxidation in the presence of antibodies. IL-33^ox^ levels were detected using IL-33^ox^ ELISA (mean ± SD of duplicate determinations). **(e)** Impact of recombinant IL-33^red^, oxidised for 24 hours in the presence of test antibodies, on human lung epithelial cell migration. The data shown (mean ± SD of triplicate determinations) are a representative example from two independent experiments. **(f)** A schematic of the proposed mode of action of tozorakimab. IL-33^red^, that is rapidly released following epithelial damage, binds tozorakimab with a high affinity and a fast association rate. Tozorakimab directly neutralizes IL-33^red^ inflammatory activities at the ST2 receptor. The IL-33^red^–tozorakimab complex prevents the oxidation of IL-33, IL-33^ox^– RAGE/EGFR signalling and epithelial dysfunction. EGFR, epidermal growth factor receptor; ELISA, enzyme-linked immunosorbent assay; IL, interleukin; IL-1RAP, interleukin-1 receptor accessory protein; mAb, monoclonal antibody; ox, oxidized; RAGE, receptor for advanced glycation end products; red, reduced; RU, response units; SD, standard deviation.

To investigate the mechanism through which tozorakimab inhibits IL-33^ox^–RAGE-EGFR signalling, surface plasmon resonance was used to compare tozorakimab binding with IL-33^ox^ and IL-33^red^. Interestingly, the tozorakimab Fab fragment showed significantly reduced binding to immobilized IL-33^ox^ relative to IL-33^red^ with an affinity 1 400 000-fold weaker for IL-33^ox^ than for IL-33^red^ (K_D_, IL-33^ox^ 41 nM [**Fig. 6b**] vs IL-33^red^ 30 fM [**Fig. 4a**]). Consistent with the weaker binding, tozorakimab was unable to inhibit the reduced epithelial migration caused by purified recombinant IL-33^ox^ (residues = 112–270) (**Fig 6c**).

Given tozorakimab inhibition of IL-33^ox^ activity was unlikely to be mediated via direct binding, we explored whether tozorakimab could modulate the oxidation of IL-33. Tozorakimab inhibited the formation of IL-33^ox^ over 24 h (**Fig. 6d**) and restored RAGE-dependent and ST2-independent epithelial migration (**Fig. 6e**). In summary, we propose a dual mechanism of action for tozorakimab (**Fig. 6f)**. Rapid binding to IL-33^red^ inhibits the interaction with ST2 and also prevents the formation of IL-33^ox^, hence preventing downstream IL-33^ox^–RAGE/EGFR signalling.

## Discussion

Here we describe the identification of tozorakimab, a novel high affinity IL-33 antibody that is the first anti-IL-33 antibody reported to inhibit signalling through both the IL-33^red^–ST2 and IL-33^ox^–RAGE-EGFR signalling pathways. The mechanism of action of tozorakimab is therefore distinct from anti-ST2 therapeutics that only target IL-33^red^–ST2 signalling [11,31]. Tozorakimab has a superior affinity and a similar fast association rate for IL-33^red^ compared with sST2, the endogenous soluble antagonist of IL-33.

It was hypothesized that a therapeutically effective anti-IL-33 antibody would require an affinity superior to that of IL-33 for the ST2 receptor. Previous estimates of the affinity of the IL-33–sST2 interaction have varied by at least three orders of magnitude [19,21,43–46]. Therefore, we performed novel high-sensitivity KinExAs to reassess the affinity and kinetics of the IL-33–sST2 interaction in free solution. We demonstrated that the IL-33–sST2 binding interaction (K_D_ 0.09 pM) is among the highest affinity measurements to be reported [47,48]. Differences in the affinity measurements were likely to be due to different assay conditions (e.g. the immobilisation of sST2 [19,21,44–46], pH, or binding of IL-1 receptor-accessory protein to the IL-33-ST2 binary complex [43,44,49]) or the sensitivity limits of the techniques.

Characterization of the IL-33–sST2 interaction identified that association rate kinetics (K_a_, 1.5 × 10^8^ M^−1^ s^−1^) are a major contributor to this high affinity interaction. This high affinity and fast association rate are consistent with a strong electrostatic complementarity interaction between IL-33 and ST2 [44,45,50,51]. *In silico* modelling, using measurements of the kinetics of IL-33 release in vivo [4,18,21] and refined IL-33–sST2 binding parameters, also highlighted the importance of a greater than 10^7^ M^−1^ s^−1^ association rate within the affinity profile to inhibit alarmin activity. In summary, studies of the IL-33–sST2 interaction highlighted the optimal pharmacological and kinetic profile for an anti-IL-33 therapeutic antibody.

We utilized our understanding of the conformational changes associated with IL-33 oxidation [21] to devise a novel fit-for-purpose antibody selection and screening campaign using both a wild-type IL-33 (IL-33^red^) and an oxidation resistant form (IL-33^C>S^) of IL-33. Interestingly, the only antibody that fully inhibited IL-33-dependent cellular NF-κB-signalling and cytokine release was derived from selection with IL-33^C>S^. Indeed, an initial screening campaign to isolate antibodies that inhibited both NF-κB-signalling and cytokine release using IL-33^red^ was unsuccessful. It is therefore possible that utilization of a stabilized form of IL-33 contributed to our success in identification of fully neutralising antibodies by minimizing conformational changes in relevant epitopes. Even so, such antibodies appeared to be a rare occurrence in our phage libraries.

Affinity optimization (>100,000-fold) to explore antibody sequence modifications across CDR1–3 led to the identification of tozorakimab, which had a high affinity for IL-33 (K_D,_ 30 fM). Tozorakimab exceeded the affinity of sST2 for IL-33, which makes it one of the highest affinity antibodies described to date [47,52–55]. Tozorakimab had an *in vitro* potency greater than that of sST2. It fully inhibited ST2-dependent cytokine release driven by an endogenous mixture of full-length and N-terminally processed forms of IL-33^red^, across a range of primary human cells and *in vivo*. Importantly, tozorakimab neutralized IL-33 effectively in a murine model of acute lung injury characterized by the rapid release of IL-33 [18,21]. This indicated its potential to effectively overcome spikes of released IL-33 associated with disease exacerbations [36].

Oxidation of IL-33 limits its ST2-dependent activities [21]; however, recent data has demonstrated that IL-33^ox^ can signal via RAGE-EGFR on airway epithelial cells *in vitro* and induces a COPD-associated epithelial phenotype [22]. Consistent with the large structural difference between IL-33^red^ and IL-33^ox^ [21], we showed in the present study that tozorakimab is unable to directly neutralize IL-33^ox^. However, our investigations indicated that tozorakimab potently inhibited IL-33^ox^-driven RAGE/EGFR signalling via prolonged binding of IL-33^red^. This prevented its conversion to IL-33^ox^ and thereby reduced IL-33^ox^-dependent activities in epithelial cells. Potential roles for IL-33^ox^ signalling in other cell types, beyond epithelial cells, remain to be elucidated and may identify novel therapeutic opportunities for tozorakimab as a treatment for human disease.

Tozorakimab has completed phase 1 clinical evaluation, in which it was shown to be well-tolerated with no safety concerns in healthy subjects and patients with mild COPD (NCT03096795) [56,57]. Building on the pre-clinical pharmacology studies described here, pharmacodynamic biomarker analyses from a phase 1 study suggest that tozorakimab reduced biomarkers of inflammation, including serum IL-5, IL-13 and blood eosinophils in patients with mild COPD [57]. Tozorakimab is currently being investigated in multiple inflammatory diseases including a phase 3 study in acute respiratory failure (NCT05624450); phase 2 (NCT04631016) and phase 3 (NCT05166889 and NCT05158387) trials in COPD and phase 2 studies in asthma (NCT04570657) and diabetic kidney disease (NCT04170543).

## Methods

### Protein reagents

The expression and purification of the protein reagents and the biotinylation of IL-33 have been described previously [21]. Protein mass integrity [58] was assessed using sodium dodecyl sulphate–polyacrylamide gel electrophoresis (SDS-PAGE) and mass spectrometry under reduced and non-reduced conditions. Receptor glycoproteins and Fab fragments were analysed and re-monomerized using Superdex 200 size exclusion chromatography (SEC). Protein concentrations were assessed by ultraviolet spectrophotometric methods. Further details can be found in the Supplementary methods.

### KinExA

KinExA measurements were performed on a KinExA 3200 (Sapidyne Instruments, Boise, Idaho, USA) instrument and data were processed using the KinExA Pro software version 4.3.10 (Sapidyne Instruments, Boise, Idaho, USA). Details regarding the preparation of the sST2–IL-33 and tozorakimab–IL-33 equilibration mixtures, sample handling, sampling bead column preparation and secondary detection conditions for each experimental system can be found in the **Supplementary methods**.

### Kinetic modelling of IL-33 antibody binding

*In silico* modelling was used to emulate the *in vivo* suppressive effect of neutralizing antibodies (150 mg every 4 weeks) with varying binding properties on circulating IL-33 levels in the healthy state (steady-state, without drug) and post dose (**Supplementary Fig 1a)**. Detailed methods are provided in the **Supplementary methods**.

### Antibody binding characterization by surface plasmon resonance

Biosensor affinity and kinetic assessment experiments were performed at 25°C on Biacore 2000 or 8K instruments (Cytiva, Little Chalfont UK) using chips and reagents from Cytiva (Little Chalfont, UK). The running buffers used throughout were based on HEPES (4-(2-hydroxyethyl)-1-piperazineethanesulfonic acid)-buffered saline-based buffer concentrates (HBS-EP+, Cytiva).

Primary affinity characterization of tozorakimab precursor antibodies was performed using protein G’-mediated capture immobilization of intact antibodies and flow of IL-33^red^. The preparation of protein G’ surfaces on Biacore CM5 chips (Cytiva) has been described previously [41,59,60] and is reported briefly in the **Supplementary methods**. Binding of recombinant Fab to immobilized biotinylated IL-33 variants was assessed using Biacore 8K instruments as described in the Supplementary methods.

### Isolation of lead antibodies for IL-33 from a naive phage display library

Large libraries of scFv human antibody, which were based on variable genes isolated from B-cells from adult naive donors and cloned into a phagemid vector based on the filamentous phage M13 [37,38], were used for selection. IL-33-specific scFv antibodies were isolated from the phage display library in a series of repeated selection cycles using recombinant human IL-33^C>S^ and IL-33^red^ [39] (brief details can be found in the **Supplementary methods)**.

### Affinity maturation through phage display selection

A targeted mutagenesis approach was used for affinity maturation of the germline version of the lead mAb using large scFv-phage libraries created by oligonucleotide-directed mutagenesis of the CDRs [61]. The libraries were subjected to affinity-based phage display selection to select variants with a higher affinity for human IL-33^red^ than the lead mAb. Generally, the selection and phage rescue were performed as described for the naive phage library (**see Supplementary methods**). There were five selection cycles in the presence of decreasing concentrations of biotinylated antigen (typically 50 nM–10 pM).

### Receptor–ligand competition assays

A homogeneous fluorescence resonance energy transfer (FRET) HTRF® (Homogeneous Time-Resolved Fluorescence, Cisbio International) assay was used to measure the inhibition of the IL-33–sST2 interaction. Please see the **Supplementary methods f**or full details.

### Epitope competition assay for the comparison between the lead mAb and tozorakimab, and for the determination of species cross-reactivity and selectivity for IL-33

Samples were tested for the inhibition of biotinylated Avi-tag histidine-tag IL-33^red^ or IL-33^C>S^ binding to anti-IL-33 IgG (33640117) or Avi-tag histidine-tag IL-33^red^ binding to tozorakimab using a similar method to the receptor–ligand assay. Please see the **Supplementary methods** for full details.

### Cell culture and *in vitro* pharmacology assays

HUVECs were maintained in EBM-2 culture media according to the manufacturer’s instructions (Lonza). NF-κB signalling and cytokine release assays have been described previously [21]. Human lung tissue was supplied in Aqix RS-I medium (Aqix Ltd) by the Royal Papworth Hospital Research Tissue Bank with written consent and study approval from the NRES East of England (Cambridge East) Research Ethics Committee (reference number 08/H0304/56 + 5) and all methods were carried out in accordance with their guidelines and regulations. Human lung tissue was homogenized in phosphate-buffered saline (PBS) for 30 seconds using a tissue homogenizer and cell debris was removed by centrifugation at 13,000 rpm [18]. A dilution of lysate that stimulated a half-maximal IL-8 release was selected for antibody neutralization studies.

The collection and culture of human blood-derived mast cells from three donors is described in the S**upplementary methods**. For assessment of cytokine release, IL-6 and IL-3 were removed from the culture 24 hours before stimulation with IL-33^C>S^. Human blood-derived mast cells were plated between 40,000–50,000/well in StemSpan (StemCell Technologies) serum-free medium supplemented with Stem Cell Factor (Peprotech) 100 ng/mL and IL-9 (Peprotech) 20 ng/mL. Antibodies were evaluated against a final concentration of IL-33^C>S^ at 0.3 ng/mL. Cells were incubated overnight, and supernatants were analysed using the MSD (MesoScale Discovery) U-plex Biomarker Group 1 human Multiplex assay after 20–24 hours.

Human peripheral blood mononuclear cells (PBMCs) were sourced from healthy donors who had provided informed consent, with ethical approval from the Research Tissue Bank (RTB; ethics number RTB, 16/EE/0334) and all methods were carried out in accordance with their guidelines and regulations. The collection and culture of PBMCs is described in the **Supplementary methods**. PBMCs were stimulated with human IL-12 (5 ng/mL) and IL-33^C>S^ (0.1 ng/mL). Cells were incubated at 37°C, 5% CO_2_ for 48 hours. Media supernatants were collected and soluble IFN-γ was measured by enzyme-linked immunosorbent assay (ELISA) using anti-human IFN-γ capture antibody (Pharmingen, 551221), biotinylated anti-human IFN-γ detection antibody (Pharmingen, 554550) and DELFIA Europium-labelled streptavidin (PerkinElmer), for fluorescent detection and quantification.

Data were analysed using Prism 8 or Prism 9 software (GraphPad). IC_50_ values and the 95% CI were determined by curve fitting using a three or four parameter logistic equation or log (inhibitor) vs the normalized response with variable slope.

### Biochemical IL-33 oxidation assays

IL-33^red^ (10 µg/mL) was incubated in Iscove’s modified Dulbecco’s medium with 1% bovine serum albumin (BSA) at 37°C in the presence or absence of tozorakimab, control mAb or lead mAb. At time 0, 0.5, 1, 2, 3, 4, 6, 8, 9 and 24 hours) 10 µL was removed, diluted to 1 µg/mL in PBS/1% BSA, snap frozen on dry ice and stored at – 80°C. Samples were analysed using the human IL-33 duoset (R&D Systems) and the IL33004-IL330425-biotin assays (AstraZeneca), detailed in [21] for measuring IL-33^ox^ and IL-33^red^, respectively.

### Epithelial scratch wound repair assays

A549 cells (ATCC, CCL-185) were cultured in RPMI medium with GlutaMAX (Thermo Fisher Scientific Inc., 61870036) containing 10% fetal bovine serum (FBS) and 1% penicillin/streptomycin. Cells were harvested using accutase (PAA Laboratories, L11-007), seeded at 5 × 10^5^ cells/100 μL into Incucyte Imagelock 96-well plates (Sartorius, BA-04857) and incubated for 6 hours. Cells were washed, serum starved (RPMI medium with GlutaMax without FBS) and incubated for 16–20 h. Cells were scratched using a WoundMaker (Essen Bioscience), washed and treated with RPMI medium with GlutaMAX supplemented with 0.1% (v/v) FBS and 1% (v/v) penicillin/streptomycin. Cells were incubated with 0.001–67 nM tozorakimab, a human IgG1 isotype control (67 nM) or were left untreated. Incucyte S3 systems (Essen Bioscience) were used for wound closure imaging and analysis over a 72-h period. Relative wound density was calculated using the wound closure algorithm (Essen Bioscience) and Incucyte S3 software (Essen Bioscience).

### Lung epithelial cell migration assay

A549 cells were cultured in F12K Media (Gibco) containing 10% FBS and 1% penicillin/streptomycin. Cells were harvested with 0.5% Trypsin-ethylenediaminetetraacetic acid (EDTA, Gibco), seeded at 1 × 10^5^ cells/well in 96-well plates and incubated for 24 hours. The cells were serum starved (F12K without FBS) for 24 hours. To test the effect of the antibodies, 1 nM tozorakimab, anti-ST2 (WO 2013/173761 Ab2; SEQ ID 85 and SEQ ID 19), anti-RAGE m4F4 (WO 2008137552) and isotype control antibodies were combined with IL-33^ox^ and incubated with the cells for 24 hours before the assessment of cell migration. Alternatively, antibodies were combined with IL-33^red^ in Iscoves’s Modified Dulbecco’s Medium (IMDM, Gibco) overnight before the treatment of the cells for 24 hours.

Following treatment, cell migration was evaluated using a 96-well Transwell system (Corning, CLS3422-48EA, 8 mM pore size). F12K media containing 10% FBS was added to the bottom chamber of the Transwell system. Treated cells were harvested with trypsin, resuspended in F12K media without FBS and added to the top chamber of the Transwell system. After 16 hours, cells that had migrated into the bottom chamber were quantified using Cell Titer Glo (Promega Corporation) according to the manufacturer’s instructions. Plates were incubated for 10 min at room temperature and luminescence was then acquired on a Cytation 5 (BioTek) with an integration time of 0.2 seconds.

### *Alternaria alternata* allergen challenge model

All in vivo work was carried out to UK Home Office ethical and husbandry standards under the authority of an appropriate Project Licence (which was approved by Babraham Institute Animal Welfare and Ethical Review Body (AWERB) and aligned to the ARRIVE guidelines. The effects of tozorakimab were assessed in a humanized IL-33 mouse model of *A. alternata* induced airway inflammation [21]. Female humanized IL-33 mice (6–10 weeks of age) were anaesthetized using isoflurane. Either 25 µg of *A. alternata* extract (Greer, Lenoir, NC) or PBS vehicle was administered intranasally (volume, 50 µL). Mice received tozorakimab (10, 3, 1, 0.1 mg/kg), NIP228 isotype control IgG (10mg/kg) or vehicle (PBS, 10 mg/kg) intraperitoneally, 24 hours before the intranasal *A. alternata* challenge. There were 6 mice per group, the experimental unit is the animal, and a complete randomized design was used with one intervention factor of interest (the treatment). The allocation of the animals to cages and subsequent treatment was pseudorandom at the time of unpacking and treatment order was not randomized. Power analysis was performed using previously obtained bronchoalveolar lavage IL-5 measurements from the same model (mean 125 ng/mL, standard deviation = 63) and the formula n = (Za + Zb)^2^ × (2s^2^)/(m1 − m2)^2^. Using a power (beta) of 80% and alpha = =0.05, it was determined that using n = 6 would enable us to see a 72% change from the control assuming the variance in the present study was identical. Mice were terminally anaesthetized using pentobarbital sodium (400mg/kg intraperitoneal) 24 h after the challenge, followed by exsanguination. Tracheal cannulation was then performed to enable bronchoalveolar lavage. BALF was collected using PBS 1 mL (sequential lavage with 0.3, 0.3 and 0.4 mL volumes), followed by centrifugation and the supernatant was stored at –80°C before the analysis of IL-5 using ELISA (Meso Scale Discovery). The analysis was conducted by the scientist collecting the data and was not blinded. Measurement of the tozorakimab serum concentration by ELISA in the treated groups identified three animals that had been mis-dosed with a serum concentration below the lower limit of quantification. These were excluded from the analysis.

## Acknowledgements

We thank Laura Drought, PhD, and Richard Claes, PhD, from PharmaGenesis London, London, UK, who provided medical writing support, which was funded by AstraZeneca. We would also like to thank Delphine Fougeron, Feenagh Keyes and Trevor Wilkinson for scientific input and Natasha Karp for statistical support.

## Author contributions

EE, DGR, ICS, DTYC, MP, RDM, KAV, RJB, TM, DCL, CC and ESC, contributed to the concept and design of the study. EE, DGR, ICS, SC, DTYC, CECH, KFH, JBM, LR, DAS, CH, ECH, MDS, BPK, DJC and ESC were involved in data acquisition. EE, DGR, ICS, SC, DTYC, CECH, KFH, TE, JBM, LR, DAS, CH, ECH, MDS, BPK, DJC, KAV and ESC analysed the data. EE, DGR, ICS, DTYC, TE, KAV, RJB, TM, TJV, DCL, CC and ESC interpreted the data. All authors were involved in the drafting of the manuscript, providing critical revisions for important intellectual content and approving the final version submitted for publication. All authors agreed to be accountable for all aspects of the work.

## Data availability

Data supporting the results described in this manuscript may be obtained from the corresponding author upon reasonable request.

