## Supplementary Information for "Tozorakimab (MEDI3506): a dual-pharmacology anti-IL-33 antibody that inhibits IL-33 signalling via ST2 and RAGE/EGFR to reduce inflammation and epithelial dysfunction"

^1^Biologics Engineering, R&D, AstraZeneca, Cambridge, UK. ^2^Translational Science and Experimental Medicine, Research and Early Development, Respiratory & Immunology, BioPharmaceuticals R&D, AstraZeneca, Cambridge, UK. ^3^Bioscience Asthma and Skin Immunity, Research and Early Development, Respiratory & Immunology, BioPharmaceuticals R&D, AstraZeneca, Cambridge, UK. ^4^Drug Metabolism and Pharmacokinetics, Research and Early Development, Respiratory & Immunology, BioPharmaceuticals R&D, AstraZeneca, Gothenburg, Sweden. ^5^Early Oncology DMPK, Oncology R&D, AstraZeneca, Cambridge, UK. ^6^Bioscience Asthma and Skin Immunity, Research and Early Development, Respiratory & Immunology, BioPharmaceuticals R&D, AstraZeneca, Gaithersburg, MD, USA. ^7^Bioscience In Vivo, Research and Early Development, Respiratory & Immunology, BioPharmaceuticals R&D, AstraZeneca, Cambridge, UK.

^8^Division of Rheumatology, Department of Medicine, University of Washington, Seattle WA, USA

*Corresponding author

^a^These authors contributed equally.

### Supplementary methods

#### **Protein reagent quality control.** Superdex 200 Increase 10/300 columns and PD-10 desalting columns were supplied by Cytiva (Little Chalfont, UK). BioSep-Size Exclusion Chromatography (SEC)-S2000 columns (300 × 7.8 mm) were from Phenomenex, (Macclesfield, UK). SEC was run on an Agilent HP1100 HPLC system at 0.5 mL min^-1^ flow rate. The Amicon Ultra-4, 10 000 molecular weight cut-off centrifugal concentrators were from Millipore (Watford, UK). Sodium dodecyl sulphate-polyacrylamide gel electrophoresis (SDS-PAGE) was performed using 4–20% 1.5 mm tris-glycine gels and the supporting reagents were from Life Technologies Limited (Paisley, UK).

All proteins were assessed for mass integrity using SDS-PAGE and mass spectrometry (MS) under reduced and non-reduced conditions [1]. Glycoprotein masses were assessed by matrix-assisted laser desorption/ionization-time-of-flight MS in the positive ion mode using a ProteinChip Reader PBS II instrument and ProteinChip software version 3.2.1 (Seldi Service Ltd, Kingston upon Thames, UK, formerly manufactured by Ciphergen, then Bio-Rad) using NP-20 chips and EAM-1 matrices (Bio-Rad). Samples for matrix-assisted laser desorption/ionization-time-of-flight MS analyses were prepared by diluting protein samples with a saturated EAM-1 matrix solution of acetonitrile:water:trifluoroacetic acid (50:50:0.5). Receptor glycoproteins and antigen-binding fragment (Fab) were also analysed and
re-monomerized using Superdex 200 SEC. Interleukin (IL-33) forms were analysed via liquid chromatography electrospray ionization MS (Waters Xevo G2-XS) and analytical SEC. The key reagent, tag-free human IL-33 (residues 112–270), had a predicted wild-type (WT) mass of 17,994.1 Da in the fully reduced state, and the liquid chromatography-MS confirmed mass was 17,994 Da under non-reducing conditions.

Protein concentrations were assessed by ultraviolet spectrophotometric methods using a 280 nm extinction coefficient calculated from the complete amino acid sequence of the protein construct (post-translational modification-related mass changes were not included in the concentration calculations) [2].

#### **Kinetic exclusion assay (KinExA).** Soluble-serum-stimulated 2 (sST2)–IL-33 and tozorakimab–IL-33 equilibration mixtures were prepared in KinExA sample buffer (KSB) comprising Dulbecco’s phosphate-buffered saline (DPBS; Gibco catalogue number 14190144) and supplemented with 1 mg mL^-1^ bovine serum albumin (BSA) and 0.02% sodium azide. The instrument running buffer was KSB prepared without BSA. Owing to the long equilibration times, all buffers used in the KinExA experiments were sterilized using a 0.2 μm filter.

For the sampling bead columns, dry UltraLink Biosupport azlactone beads (Thermo Fisher Scientific Inc. 53110) were mixed with appropriate quantities of the reduced IL-33 (IL-33^red^) antigen or streptavidin (Thermo Fisher Scientific Inc. 21125) in 50 mM sodium hydrogen carbonate (pH 8.4) at room temperature with constant agitation for around 1.5–2 hours. Rinsing/bead-blocking steps were then performed with a solution of 10 mg mL^-1^ BSA in 1 M Tris pH 8.7. Before use, the bead slurry was resuspended in filtered KSB. The mono-biotinylation of IL-33^red^ via a reactive thiol has been described in Cohen, E.S. *et al.* 2015 [3].

Based on the reagents in the equilibration mixtures, the fluorescent secondary detection reagent used were either Alexa Fluor 647-labelled goat anti-human fragment crystallizable (Fc) antibody (Jackson Immunoresearch, 109-605-098) for the whole immunoglobulin G (IgG) assay or DyLight 650-labelled anti-His6 specific mouse monoclonal antibody (Abcam, Cambridge, UK, ab117504) for the monomerized Human Embryonic Kidney Epstein–Barr virus nuclear antigen (HEK-EBNA) expressed N-terminal–Flag-His10-tag human-sST2 glycoprotein (monomeric sST2, 19-328 extracellular domain, Uniprot Q01638) and the histidine (His)-tagged Fab assays. Fluorescence signal artefacts that were especially prevalent in assays using the His-tag based secondary detection reagent (Fab-His, serum-stimulated 2 [ST2]-Flag-His10 and DyLight 650-labelled anti-His6) were efficiently suppressed by 0.2 μm filtration of the diluted secondary detection mixture.

For all experiments, the equilibrating samples and the entire KinExA 3200 instrument was housed within a temperature-controlled cabinet (Series 3 HTCL 750 Temperature Applied Sciences Ltd., West Sussex, UK) at 25°C. Bead stability was enhanced by maintaining an ice bath around the bead vial throughout the analysis. The latter modification eliminated the need to invoke drift correction during data analyses.

Simultaneous (global) analysis of the individual datasets (from at least two fixed concentrations of IgG, Fab or ST2 per interaction) were performed to give a 1:1 affinity (*K*_D_) estimate (Global N-Curve Analysis, KinExA Pro 4.3.10, Sapidyne Instruments). As well as the affinity *K*_D_ estimate, these analyses provided the active constant binding partner, which is an estimate of the fraction of IgG, Fab or sST2 that was active and participating in the interaction.

#### **Surface plasmon resonance.** Antibody-binding interactions with IL-33 were assessed using surface plasmon resonance. Chip immobilization of biotinylated IL-33 variants was achieved via amine-coupled streptavidin (Thermo Fisher Scientific Inc. 21125). Whole-human antibodies were reversibly chip-captured via amine-coupled recombinant protein G’ (protein G prime, a fragment of protein G, Sigma-Aldrich, P-4689).

To evaluate whole-antibody binding to IL-33^red^, recombinant protein G’ was reconstituted in water and buffer exchanged into D-PBS via a PD-10 desalting column Cytiva (Little Chalfont, UK) to remove Tris. This buffer-exchanged protein G’ was further diluted into 10 mM sodium acetate (pH 3.6) and was amine coupled to Biacore CM5 chips Cytiva (Little Chalfont, UK). Each analysis cycle involved the titration of 100–400 response units of each antibody onto the protein G’ surface at 5 mL min^-1^, followed by the performance of IL-33 binding analysis steps at a higher 50 mL min^-1^ flow rate, with an association time of 1.5 min. Dissociation (HBS-EP+ only flow) was followed for 7 minutes. The regeneration of protein G’ surfaces was performed with two consecutive 20-second injections of 6 M guanidine hydrochloride in D-PBS. Data were analysed using the 1:1 Langmuir model and BIAevaluation 3.1 software Cytiva (Little Chalfont, UK).

Recombinant Fab binding to immobilized biotinylated IL-33 variants was performed on a Biacore 8K instrument Cytiva (Little Chalfont, UK). Streptavidin chip surfaces were prepared using lyophilized streptavidin reconstituted with DPBS. Streptavidin was diluted to 4 mg mL^−1^ in 10 mM sodium acetate (pH 4.5) and covalently immobilized to selected flow cell surfaces of a C1 chip by standard amine coupling methods. Final streptavidin surfaces in the range of 65–70 response units were achieved. N-terminally Avi-tag biotinylated IL-33 variant (BirA biotinylated tagged IL-33^red^ or an analogous tagged oxidized IL-33 variant) or minimally amine biotinylated bovine ubiquitin control protein was titrated onto the streptavidin surface. The quantity of IL-33^red^ titrated onto the streptavidin surface allowed binding of 40 response units of Fab at saturation, which represented 90% of the predicted binding response (R**_max_**). This relatively low level of analyte (Fab) binding minimized mass-transport-induced artefacts, especially when combined with relatively rapid 50 mL min^-1^ assay flow rates [4,5].

#### ***In silico* modelling methods.** The first step in the modelling process was to assess the effect of the varying IL-33 release and degradation parameters. Initial parameter values were extracted from the IL-33 time course data measured in bronchoalveolar lavage and plasma using the *Alternaria alternata* (ALT) challenge study as described in Cohen *et al.* 2015 [3]. The IL-33 release rate varied over a range of 5–120 minutes, with an assumed overall antibody affinity of 0.1 pM and an IL-33 degradation half-life set at 60 minutes to evaluate the influence of the release rate parameter (**Supplementary Fig. S1b–e**). The antibody association rate varied across four orders of magnitude (10^5^, 10^6^, 10^7^, 10^8^ M^−1^ s^−1^) to assess its effect on IL-33 spike suppression. An antibody affinity of 0.1 pM was chosen based on the sST2–IL-33^red^ affinity results (see **Fig. 1a** in the main manuscript).

We then assessed the effect of a varying IL-33 degradation half-life over 30–300 minutes with the antibody affinity set at 0.1 pM and the IL-33 release half-life at 15 minutes (**Supplementary Fig. S1f-i**). Again, the antibody association rate was varied across four orders of magnitude (10^5^, 10^6^, 10^7^, 10^8^ M^−1^ s^−1^). From these model outputs a detailed IL-33 spike suppression profile was produced (**Fig. 1c** in the main manuscript). These outputs were for four association rates for two antibody overall affinities of 10 and 0.1 pM, with the IL-33 release rate and IL-33 degradation half-life fixed at 15 minutes and 1 hour, respectively.

Please refer to the legend of Supplementary Fig S1a for additional explanation of silico modelling methods and assumptions.

#### **Isolation of lead antibodies to IL-33 from a naive phage display library.** Briefly, the single-chain variable fragment (scFv)-phage particles were incubated with biotinylated recombinant IL-33^red^ or oxidation-resistant IL-33 (IL-33^c>s^) as antigens in solution, typically for 2 hours. ScFv bound to the antigen was captured on streptavidin-coated paramagnetic beads (Dynabeads^®^, M-280). Unbound phage was removed in a series of wash cycles using PBS-Tween. Phage particles retained on the antigen were eluted, infected into bacteria and rescued for the next round of selection [6]. Typically, 2–3 rounds of selection were performed with decreasing antigen concentrations (100, 50 and 25 nM) to increase stringency. ScFvs were converted to human IgG1 as previously described [7].

#### **Receptor–ligand biochemical assays.** Samples were tested for the inhibition of biotinylated Avi-tag His-tag IL-33^red^ or IL-33^C>S^ binding to sST2-Flag His by adding 5 µL of each test sample to a 384-well low-volume assay plate (Costar, 3673). A solution (2.5 µL) containing 2.4 nM of either biotinylated Avi-tag His-tag IL-33^red^ or IL-33^C>S^ pre-combined with 6 nM streptavidin Eu3+ cryptate (Cisbio International, 610SAKLB) for 1 hour was added. Then 2.5 µL of 4 nM sST2 FLAG His combined with 20 nM anti-FLAG XL665 (Cisbio International, 61FG2XLB) was added. All dilutions were performed in assay buffer containing 0.8 M potassium fluoride (VWR, 26820.236) and 0.1 % BSA (Sigma, A9576) in DPBS (Invitrogen, 14190185). Assay plates were incubated for 1 hour at room temperature, followed by 19 hours at 4°C. Time-resolved fluorescence was read using a 320 nm excitation filter and 620 nm and 665 nm emission filters (100 flashes) and an EnVision plate reader (PerkinElmer).

Results were calculated from the 665/620 nm ratio and expressed as percentage of DELTA F (sample ratio − negative control ratio/negative control ratio × 100). The negative control was defined by the absence of biotinylated Avi-tag His-tag IL-33^red^ or IL-33^C>S^. Data were expressed as percentage of specific binding ([sample − non-specific binding (NSB)/total −NSB] × 100) and analysed using a four-parameter logistic equation with GraphPad Prism 6.01 (GraphPad Inc., California) to give half maximal inhibitory concentration(IC_50)_ values: Y = bottom + (top − bottom)/(1 + 10^([LogIC50 − X] × HillSlope])^) where X is the logarithm of concentration and Y is the percentage of specific binding.

#### **Epitope competition assays.** Test samples (5 µL) were added to a 384-well low-volume assay plate (Costar, 3673). A solution (2.5 µL) containing 0.12 nM of either biotinylated Avi-tag His-tag IL-33^red^ or IL-33^C>S^ pre-combined with 0.75 nM streptavidin Eu3+ cryptate (Cisbio International, 610SAKLB) was added. Then, 2.5 µL of 10 nM 33640117 [8] human IgG1 labelled with DyLight 650 was added. All dilutions were performed in assay buffer containing 0.8 M potassium fluoride (VWR, 26820.236) and 0.1% BSA (Sigma, A9576) in DPBS (Invitrogen, 14190185). Assay plates were incubated for 1 hour at room temperature, then 19 hours at 4°C. Time-resolved fluorescence was read using a 320 nm excitation filter and 620 nm and 665 nm emission filters (100 flashes) and an EnVision plate reader (PerkinElmer). The results were calculated using the same method as the receptor–ligand assay.

#### **Cell culture.** ***Preparation of human blood-derived mast cells.*** Human blood-derived mast cells were collected from three donors. Briefly, peripheral blood mononuclear cells (PBMCs) were collected from a leukopak (ALLCells) and mixed with Robo II Buffer (StemCell Technologies) at a ratio of 1:4 and centrifuged at 500 x g for 15 minutes. The pellet was resuspended in 1X Vitalyse (CMG) for red blood cell lysis and incubated for 30 minutes. Cells were further washed with Robo II Buffer twice. PBMCs were then progenitor enriched (StemCell Technologies). Enriched cells (30 – 60 ×10^6^) were stained with anti-CD34 (Clone 581) FITC, anti-CD1117(Clone 104D2) PE and anti-FcER1a (CRA-1) APC (Biolegend). Cells were then sorted into a CD117+FcER1a+CD34+ population using a FACSAria Fusion Flow Cytometer (Becton Dickinson). Sorted mast cell progenitor cells were cultured in serum-free medium (StemSpan, StemCell Technologies) supplemented with stem cell factor 100 ng/mL, IL-6 100 ng/mL, IL-3 30 ng/mL and IL-9 20 ng/mL (Peprotech). Cell culture and expansion were continued for a minimum of 4 weeks with fresh media supplementation 1–3 times per week before use.

#### ***Preparation of PBMCs.*** Human PBMCs were sourced from healthy donors who had previously provided informed consent, with ethical approval from the Research Tissue Bank (RTB; ethics number RTB, 16/EE/0334). PBMCs were isolated from leukocyte cones using a polysucrose (Ficoll) gradient and maintained in RPMI 1640 (supplemented with 10% v/v heat-inactivated fetal bovine serum and 1% v/v penicillin/streptomycin) at 37°C in a 5% carbon dioxide, humidified atmosphere.

### Tables and figures

#### **Supplementary table S1.** KinExA-based *K*_D_ and *k*_a_ determinations with human recombinant IL-33^red^.

| **Receptor** | ***K*_D_, pM (95% CI)** | ***k*_a_, M^-1^ s^-1^ (95% CI)** |
| --- | --- | --- |
| Tozorakimab IgG mAb | 0.03 (0.008–0.062) | ND |
| Tozorakimab Fab | 0.02 (0.004–0.036) | 8.5 × 10^7^ (7.8–9.3 × 10^7^) |
| Monovalent ST2 | 0.09 (0.05–0.15) | 1.5 × 10^8^ (1.4–1.6 × 10^8^) |

CI, confidence interval; Fab, antigen-binding fragment; IgG, immunoglobulin G; IL, interleukin; *k*_a_, association rate; *K*_D_, affinity rate; KinExA, kinetic exclusion assay; mAb, monoclonal antibody; ND, not determined; red, reduced; ST2, serum-stimulated 2.

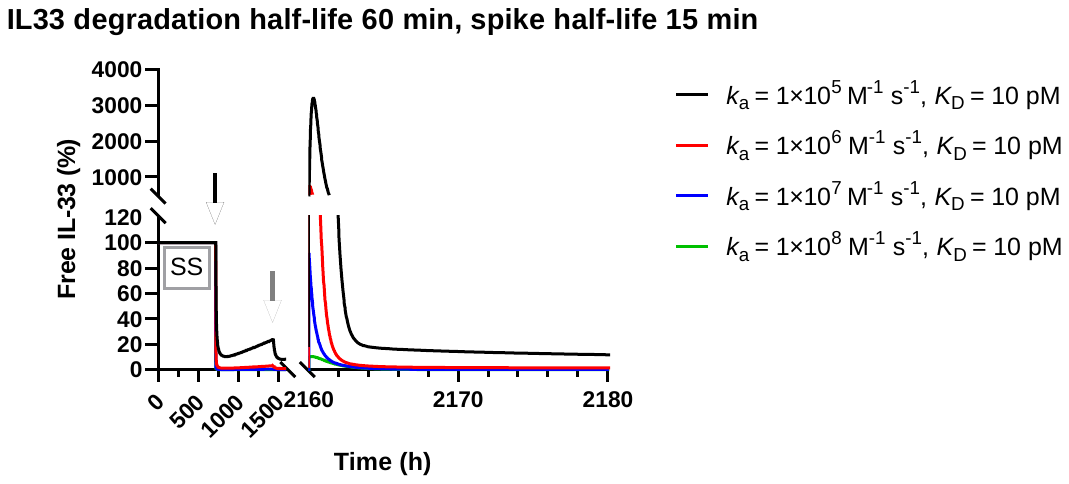

a

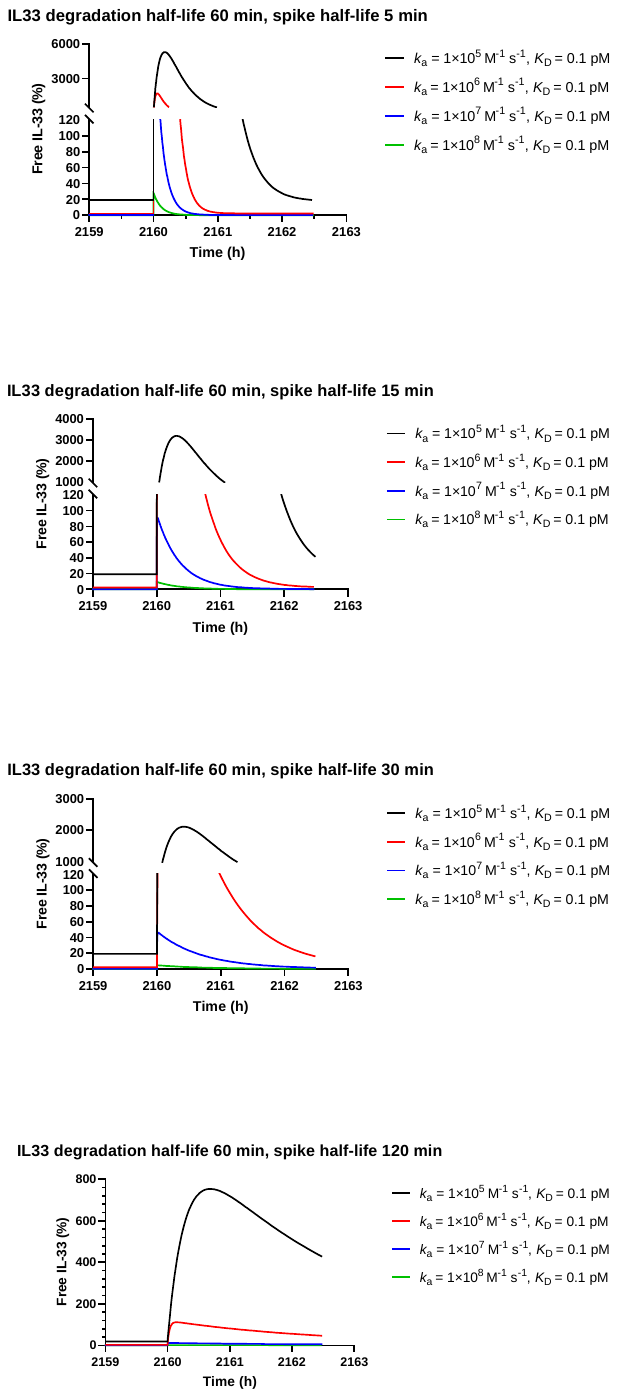

e

d

b

c

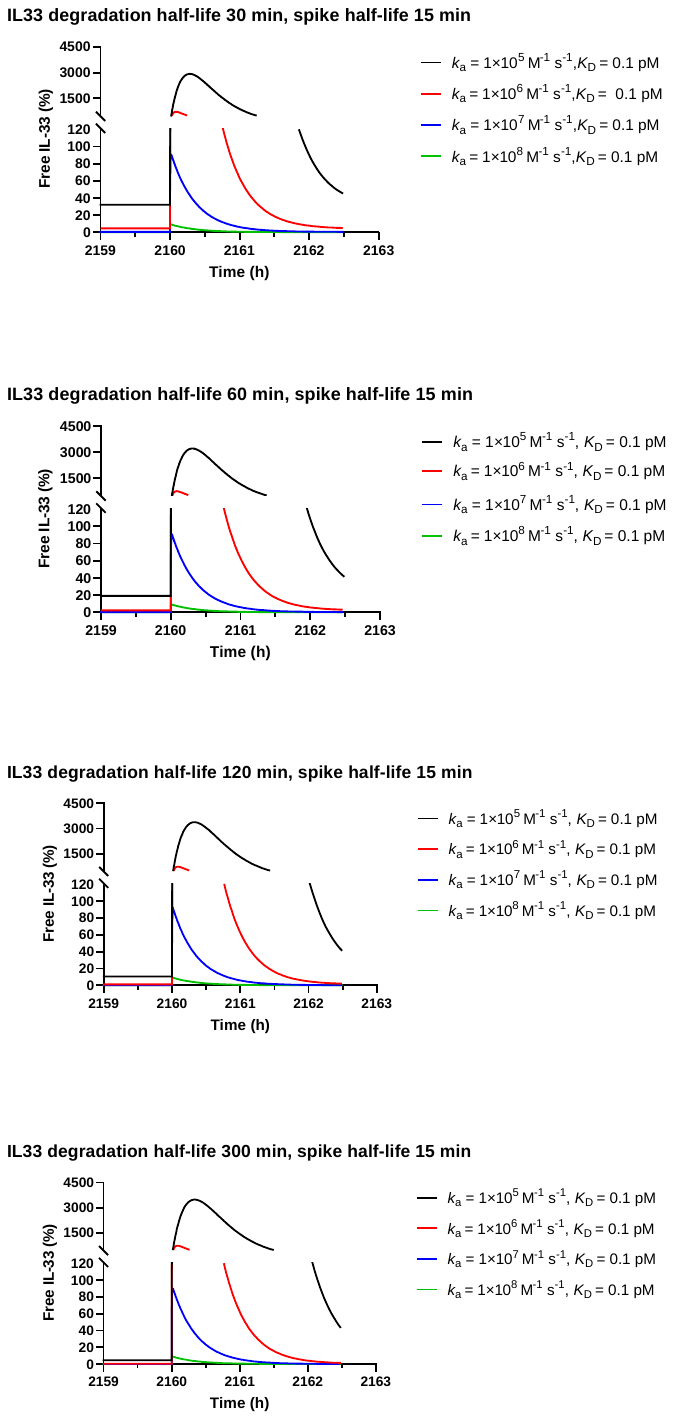

i

h

g

f

#### **Supplementary figure S1**. *In silico* modelling of the suppression profile expected from antibodies with varying kinetic and affinity profiles. (**a**) Full duration time course of the model covering all stages of the dosing regimen. The circulating steady-state IL-33 levels before the first dose of tozorakimab at 1 month (720 hours, red arrow) was set at 100%. The first monthly dosing of tozorakimab at 150 mg is indicated by the red arrow. The initial rapid IL-33 suppression profile slowly reversed until the second monthly dosing denoted by the blue arrow. At month 3, a spike of IL-33 parameterized from the ALT *in vivo* mouse model data was triggered in the *in silico* model concurrent with antibody dosing. The effects of the four *k*_a_, which varied in 10-fold steps between 10^5^ and 10^8^ M^−1^ s^−1^ range (black, red, blue and green, respectively), were predicted. The antibody *K*_D_ was set at 10 pM, the IL-33 spike half-life at 15 minutes and the IL-33 degradation rate was fixed at 1 hour. (**b–i**) Preliminary parameter range-finding examples were performed to optimize the relevance of the final model. (**b–e**) Stacked panels show a zoom into the IL-33 exacerbation spike region of the model. Predictions were based on the IL-33 spike half-life being varied over 5 minutes–120 minutes (**b** 5 minutes, **c** 15 minutes, **d** 30 minutes, **e** 120 minutes). The overall IL-33 degradation rate was fixed at 1 hour and the antibody affinity at 0.1 pM. (**f–i**) A zoom into the IL-33 exacerbation spike region of the model. Predictions were based on the overall IL-33 degradation rate period of 0.5–300 minutes (**f** 30 minutes, **g** 60 minutes, **h** 120 minutes, **i** 300 minutes). The IL-33 spike half-life was fixed at 15 minutes and the antibody affinity at 0.1 pM.

ALT, *Alternaria alternata*; IL, interleukin; *k*_a_, association rate; *K*_D_, affinity rate.

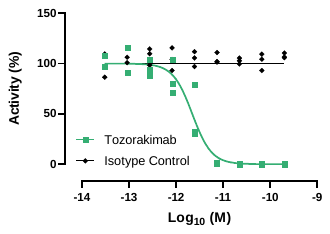

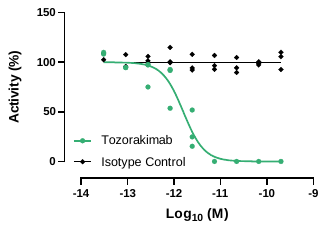
 **Mast cell IL-6 release	Mast cell IL-8 release**

b
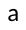

a

d

c

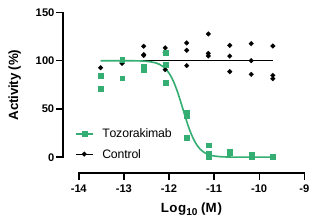

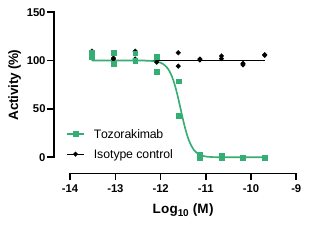
 **Mast cell GM-CSF release	Mast cell TNF-α release**

#### **Supplementary figure S2**. Effect of tozorakimab on IL-33^C>S^ (residues 112–270)-induced inflammatory responses in primary human blood-derived mast cells. (**a**) Inhibition of IL-6 release (n = 3). (**b**) Inhibition of IL-8 release (n = 3). (**c**) Inhibition of GM-CSF (n = 2). (**d**) Inhibition of TNF-α release (n = 3).

GM-CSF, granulocyte-macrophage colony-stimulating factor; IL, interleukin; TNF-α, tumour necrosis factor-alpha.

a

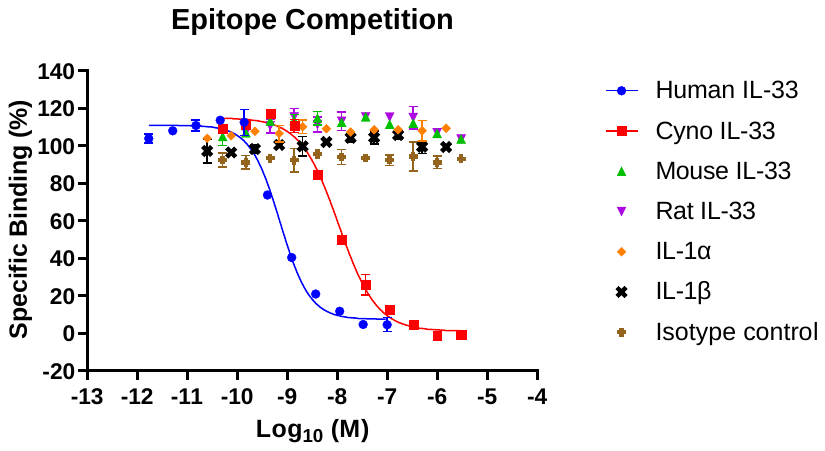

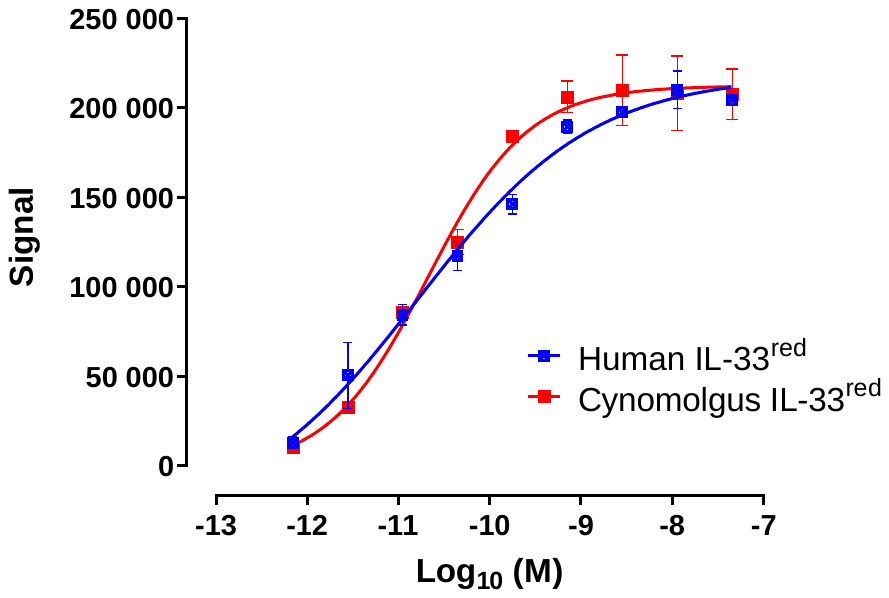

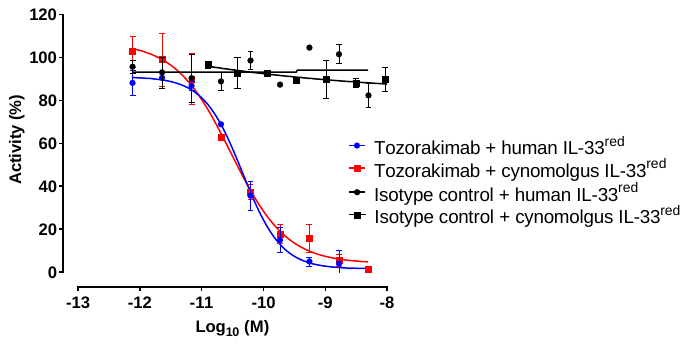
**HUVEC IL-8 release	HUVEC IL-8 release**

b

c

#### **Supplementary figure S3.** Tozorakimab selectively neutralizes human and cynomolgus monkey IL-33^red^. (**a**) Epitope competition assay for tozorakimab specificity. The ability of IL-33^red^ from cynomolgus monkey, mouse and rat, and human IL-1α and IL-1β to inhibit biotinylated Avi-tag His-tag human IL-33^red^ binding to tozorakimab (n = 2). (**b**) Recombinant human and cynomolgus monkey IL-33^red^-driven IL-8 release in HUVECs (n = 2). (**c**) Inhibition of recombinant human and cynomolgus monkey IL-33^red^-driven IL-8 release in HUVECs by tozorakimab (n = 2). Data points are presented as the mean ± SD of duplicate readings.

Cyno, cynomolgus monkey; IL, interleukin; red, reduced; His, histamine; HUVEC, human umbilical vein endothelial cells; mAb, monoclonal antibody.
